# Biodistribution of DNA-origami nanostructures in live zebrafish embryos with single-cell resolution

**DOI:** 10.1101/2023.12.22.572973

**Authors:** Christina Kolonelou, Lars Bräutigam, Steven Edwards, Enya Engström, José M. Dias, Joel Spratt, Christos Karampelias, Stefan Wennmalm, Hjalmar Brismar, Olov Andersson, Ana I. Teixeira

## Abstract

DNA origami-based nanotechnology is a versatile tool for exploring fundamental biological questions and holds significant promise for future biomedical applications. However, the development of DNA origami-based therapeutic agents is hindered by the challenge of translating *in vitro* performance into effective applications *in vivo*. Here, we exploit the optical transparency of the embryonic zebrafish to track intravenously injected, fluorescently labelled wireframe DNA origami nanostructures. Our approach integrated long-term, high-resolution imaging of transgenic live embryos with single-cell RNA sequencing, to elucidate the biodistribution of DNA nanostructures over time, up to 3 days post-injection (dpi). Notably, we observed rapid accumulation of nanostructures in the caudal hematopoietic tissue (CHT), akin to the fetal liver in mammals. We tested the effects of coating the nanostructures with an oligolysine PEG copolymer (K-PEG), a widely used strategy to enhance their stability. The K-PEG coating mitigated the accumulation rate in CHT, enabling higher percentages of the nanostructures to engage with other tissues. Additionally, our findings highlighted the pivotal role of scavenger endothelial cells in DNA origami clearance, with K-PEG offering sustained protection for the nanostructures at the CHT. Furthermore, by monitoring DNA origami in a transgenic zebrafish line designed for targeted macrophage ablation, we found that macrophages contribute to nanostructure clearance at later time points. This study introduces a framework for the analyses of the biodistribution and clearance of DNA origami nanostructures in vivo with single cell resolution and establishes a foundation for the investigation of DNA origami-based nanomedicines in animal models.

## Main

In recent years, there has been significant progress in the field of DNA nanotechnology towards enabling in vivo applications. Inherently biocompatible and biodegradable, DNA nanostructures are suitable for studies in animal models. Wireframe DNA architectures are particularly advantageous for in vivo studies as they are stable under physiological conditions due to their lower packing density compared to conventional DNA origami^1–3^. In addition, several coating approaches have been explored to enhance the stability and circulation times of DNA nanostructures in mice^4,5^. To enable tissue specific targeting, DNA origami nanostructures have been modified with antibodies, aptamers, or other bioactive ligands^6–9^. These studies have demonstrated the feasibility of using DNA origami nanostructures for targeting specific tissues, such as the immune system or tumors. In contrast, DNA nanostructures without targeting moieties have been shown to accumulate in the kidney and have been proposed as a potential therapeutic strategy to mitigate kidney damage^10,11^. Despite their promise, challenges persist in harnessing the full potential of DNA origami nanostructures as multifunctional therapeutic agents. In particular, there is a need to accurately assess the interaction profiles of DNA origami nanostructures with cells at the whole-body level as well as their dynamics of clearance. Whole-body biodistribution data acquired by imaging with positron emission tomography (PET) and bioluminescence has limitations in both resolution and sensitivity. The zebrafish (*Danio rerio*) embryo has emerged as a promising *in vivo* model in nanomedicine^12^. Its optical transparency allows for real-time observation of intravenously administered fluorescently labelled objects. Therefore, intravital imaging of zebrafish embryos can provide whole-body information on nanoparticle targeting, cellular uptake, and clearance in live animals^13–15^. In addition, zebrafish have a robust immune system resembling that of mammals^16–19^. Innate immunity starts at embryogenesis in zebrafish, marked by the emergence of macrophages from the mesoderm^20^, while adaptive immunity is activated at three weeks post fertilization^21^. This temporal separation of the onset of the innate and adaptive immunity in zebrafish embryos is advantageous for the study of the interactions of nanostructures with the immune system.

Here, we investigated the biodistribution of fluorescently labelled wireframe DNA nanostructures in live zebrafish embryos using advanced imaging methods, including light-sheet fluorescence microscopy (LSFM), Airyscan confocal microscopy, and Fluorescence Correlation Spectroscopy (FCS), coupled with single-cell RNA sequencing. Leveraging a coating strategy with K-PEG, we evaluated how surface charge modifications impact the biodistribution and clearance of DNA nanostructures. Further, we explored the role of the innate immune system using targeted ablation of macrophages in a zebrafish model. Together, our results provide insights into the performance and fate of DNA origami nanostructures in vivo.

## Results

### Wireframe NanoSheet DNA origami nanostructures for in vivo assessment

We used wireframe scaffolded DNA origami to produce single layer two-dimensional sheets (Fig. 1a, Extended Data Fig. 1a, b), herein referred to as NanoSheets (NS), which remain stable in physiological buffers, unlike DNA nanostructures constructed with tightly packed helices^22,23^. To detect the NS, we designed six protruding ssDNA strands, centrally located on their surface, which hybridize to complementary ssDNA strands conjugated with the fluorophore Texas Red (TR), referred to as NS^TR^ (Fig. 1a). Further, we employed a coating strategy widely used to enhance nanostructure stability^24^, whereby positively charged oligolysine conjugated to polyethylene glycol (K-PEG) electrostatically interacts with the negatively charged DNA nanostructures (Fig. 1a). Imaging of NS structures folded in PBS using atomic force microscopy (AFM) confirmed their self-assembly and integrity (Fig. 1b). Agarose gel electrophoresis analysis confirmed that NanoSheets folded properly and were fluorescently labeled (Extended Data Fig. 2a). Further, agarose gel electrophoresis showed that NS^TR^ coated with K-PEG, named NS^TR/K-PEG^, had a neutral surface charge and were thereby retained in the well (Fig. 1c). Removal of the K-PEG coating by incubation with chondroitin sulfate restored band migration, confirming that NS nanostructures were not structurally compromised by the coating (Extended Data Fig. 2b).

**Fig. 1:**
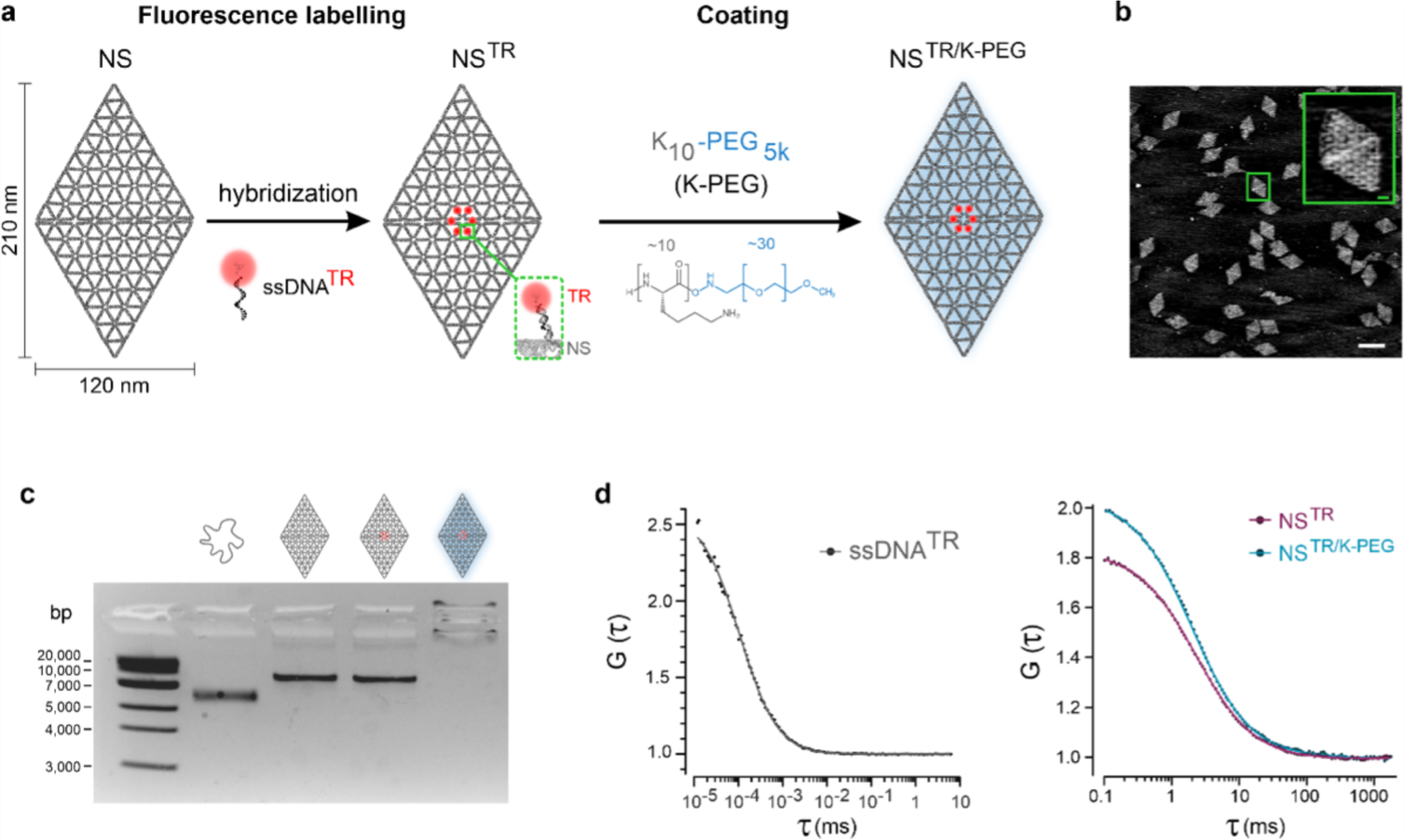
Characterization of DNA origami NanoSheets. **a.** Schematic representation of fluorescence labeling and coating strategy using Texas Red and K_10_-PEG respectively. NS: NanoSheet, NS^TR^: NanoSheet labelled with Texas Red, NS^TR/KPEG^: NanoSheet labelled with Texas Red and coated with K_10_-PEG_5K_, ssDNA^TR^: single strand DNA oligo conjugated with Texas Red. **b.** AFM phase imaging of NS nanostructures. Magnified NS image shown in the inset. Scale bar: 200 nm, scale bar inset: 20 nm. **c.** Analysis by agarose gel electrophoresis of the scaffold strand, NS, NS^TR^, and NS^TR/KPEG^ nanostructures. **d.** Characterization of ssDNA^TR^, NS^TR^ and NS^TR/KPEG^ in solution by FCS. Autocorrelation curves, where the dots represent raw data, and the solid lines correspond to two-component fit modelling. ssDNA^TR^: n= 20 technical repeats, total 6 sec traces; NS^TR^, NS^TR/KPEG^: n= 24 technical repeats each, total 5 sec traces and NS^TR/KPEG^: n= 24 technical repeats, total 5 sec traces.

Fluorescence correlation spectroscopy (FCS), a single molecule detection method, showed similar diffusion times for NS^TR^ and NS^TR/K-PEG^, indicating that the coating strategy did not induce aggregation (Fig. 1d). FCS analyses of ssDNA strands conjugated with Texas Red (ssDNA^TR^) yielded significantly shorter diffusion times compared to NS^TR^ and NS^TR/K-PEG^ (Fig1d), consistent with the lower molecular weight of ssDNA compared to NS. Importantly, the average intensity of NS^TR^ was equivalent to that of NS^TR/K-PEG^ at 7.1 kHz/molecule, indicating that the K-PEG coating did not affect the brightness of the NanoSheets. Additionally, the average intensity of ssDNA^TR^ was 3.6 kHz/molecule, suggesting that on average two molecules of Texas Red were detected per NanoSheet.

### Temporal profiles of NanoSheet distribution in zebrafish embryos

To investigate the biodistribution profiles of NS^TR^ and NS^TR/K-PEG^ in vivo, DNA nanostructures were intravenously injected into the bloodstream of zebrafish embryos at 2 days post fertilization (dpf) (Fig. 2a). We used the zebrafish model *Tg(fli1:EGFP),* where the expression of green fluorescent protein (EGFP) is under the control of the endothelial specific *fli1* promoter, enabling visualization of the vasculature of the embryo (Fig. 2b). Live embryos were imaged with LSFM for a period of 4 h, starting at 0.25 hours post injection (hpi) (Fig. 2c and Extended Data Fig. 3). We observed that NS^TR^ rapidly accumulated in the caudal hematopoietic tissue (CHT) (Extended Data Fig. 3), which is functionally equivalent to the fetal liver of mammals, responsible for embryonic hematopoiesis and the resident site of scavenger endothelial cells^25^ (Extended Data Fig. 3). At early time points, the NS^TR/K-PEG^ signal at the CHT was similar to the control (Fig. 2c). At the whole animal level, the NS^TR/K-PEG^ signal showed a different biodistribution profile compared to NS^TR^, with signal detected outside the vasculature (Extended Data Fig. 3). At 4 hpi, NS^TR/K-PEG^ mainly localized to the CHT but were also detected in other regions of the embryo, such as brain and muscle (Extended Fig.3). We confirmed that the detected fluorescence signal did not result from structurally compromised NanoSheets by injecting embryos with NS mixed with ssDNA^TR^ at a 1:1 ratio (control), which showed low fluorescence signal due to lack of retention of ssDNA in the embryo (Extended Data Fig. 3). The average fluorescence intensity of circulating NS^TR^ and NS^TR/K-PEG^ in the lumen of the dorsal aorta (DA) showed no significant differences after 1 hpi. To investigate the clearance dynamics of NanoSheets in the CHT, we measured the mean fluorescence intensity of control, NS^TR^ and NS^TR/K-PEG^ over time (Fig. 2e). The signal intensity of NS^TR^ decreased substantially up to 1 hpi and continued to decrease gradually until 4 hpi (Fig. 2c,e). In contrast, NS^TR/KPEG^ showed moderate accumulation in the CHT, with the highest fluorescence intensity detected at 2 hpi. We further investigated the fluorescence intensity profiles of NanoSheets at the CHT by confocal imaging for up to 3 dpi. We observed that the fluorescence intensity of NanoSheets decreased over time. However, only the fluorescence intensity of NS^TR^ reached control levels at 24 hpi (Fig. 2f). In addition, NS^TR/KPEG^ signal was detected at 72 hpi, which was not the case for the coating-free nanostructures. Therefore, our results suggest that the K-PEG coating resulted in pronounced stabilization of the NanoSheet fluorescence intensity.

**Fig. 2:**
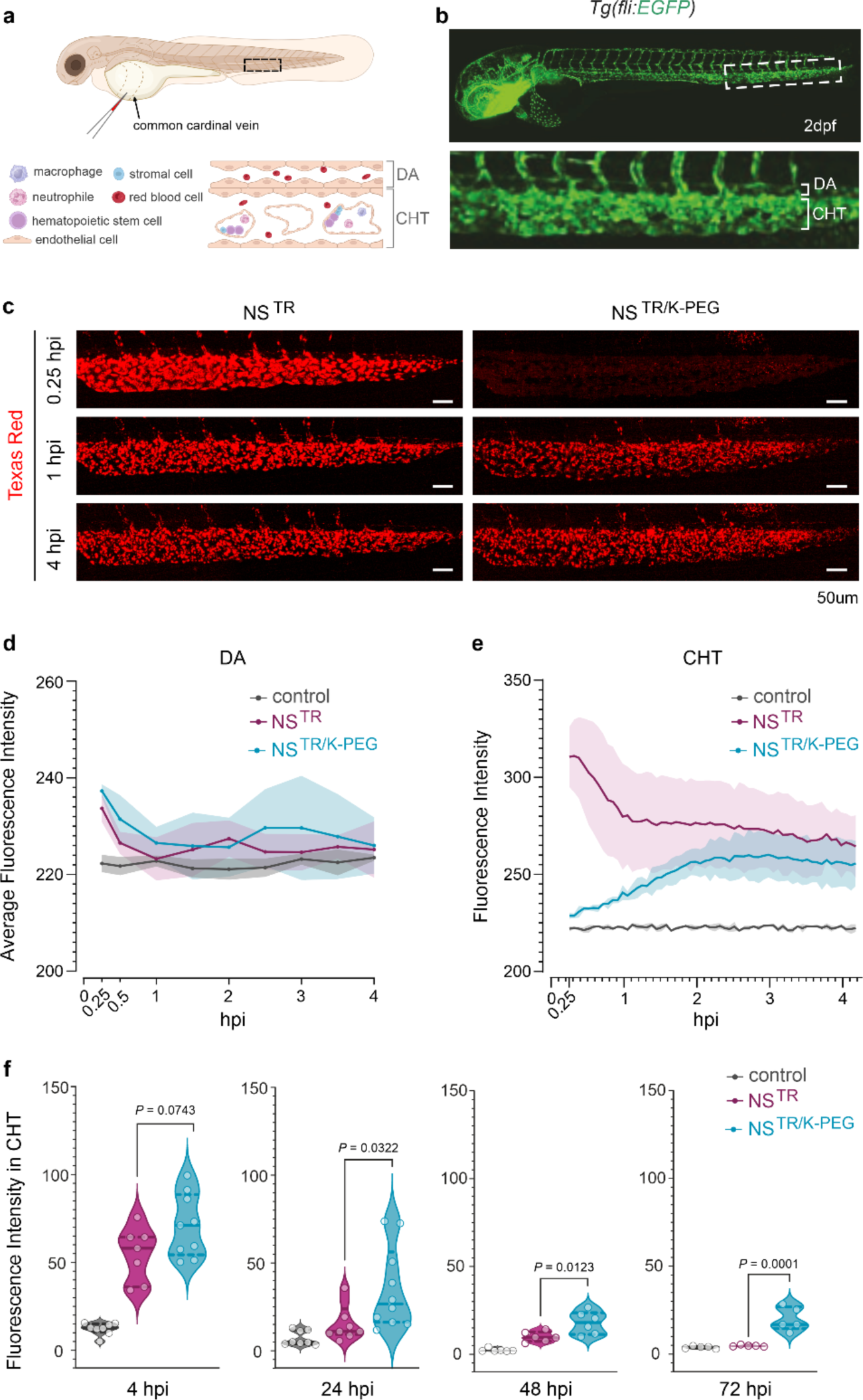
Biodistribution profiles of intravenously injected NanoSheets in zebrafish embryos. **a.** Schematic showing the site of microinjection (common cardinal vain) in a 2 day post fertilization (dpf) zebrafish embryo. Boxed region illustrating the cell types forming the dorsal aorta (DA) and caudal hematopoietic tissue (CHT) of the embryo. **b.** Whole-embryo view of a transgenic *Tg(fli1:EGFP)* zebrafish embryo, showing the vasculature (green) at 2 dpf. Magnified region corresponding to the dashed box, showing the CHT below the DA. **c.** Light-sheet fluorescence microscopy (LSFM) imaging of Texas Red signal in live embryos, showing the accumulation of the NS^TR^ and NS^TR/K-PEG^ in the CHT over time. hpi: hours post injection. Scale bar: 50 μm. **d.** Quantification of Texas Red mean fluorescence in the DA lumen from LSM imaging of embryos injected with control (NS mixed with ssDNA^TR^ in 1:1 ratio), NS^TR^ or NS^TR/K-PEG^ nanostructures at 0.25, 0.5, 1, 1.5, 2, 2.5, 3, 3.5 and 4 hpi. Values presented as average intensity from three different regions along the DA for each embryo at indicated time point (see Methods). n=3 embryos per condition. Injection of each embryo was performed as an independent experiment. Values presented as mean ± SD. **e.** Quantification of Texas Red fluorescence intensity signal from 0.25 to 4 hpi with LSM imaging at the CHT in embryos injected with control, NS^TR^ or NS^TR/K-PEG^ nanostructures. n=3 embryos per condition. Injection of each embryo was performed as an independent experiment. Values presented as mean ± SD. **f.** Long-term quantification of Texas Red signal in the CHT of live embryos by confocal imaging. Embryos were injected with control, NS^TR^ and NS^TR/K-PEG^ structures and analyzed at 4, 24, 48 and 72 hpi. n=5-10 imaged embryos per group, different embryos were imaged at each time point. *P*-values determined by one-way ANOVA followed by Tukey’s multiple comparison test.

**Fig. 3:**
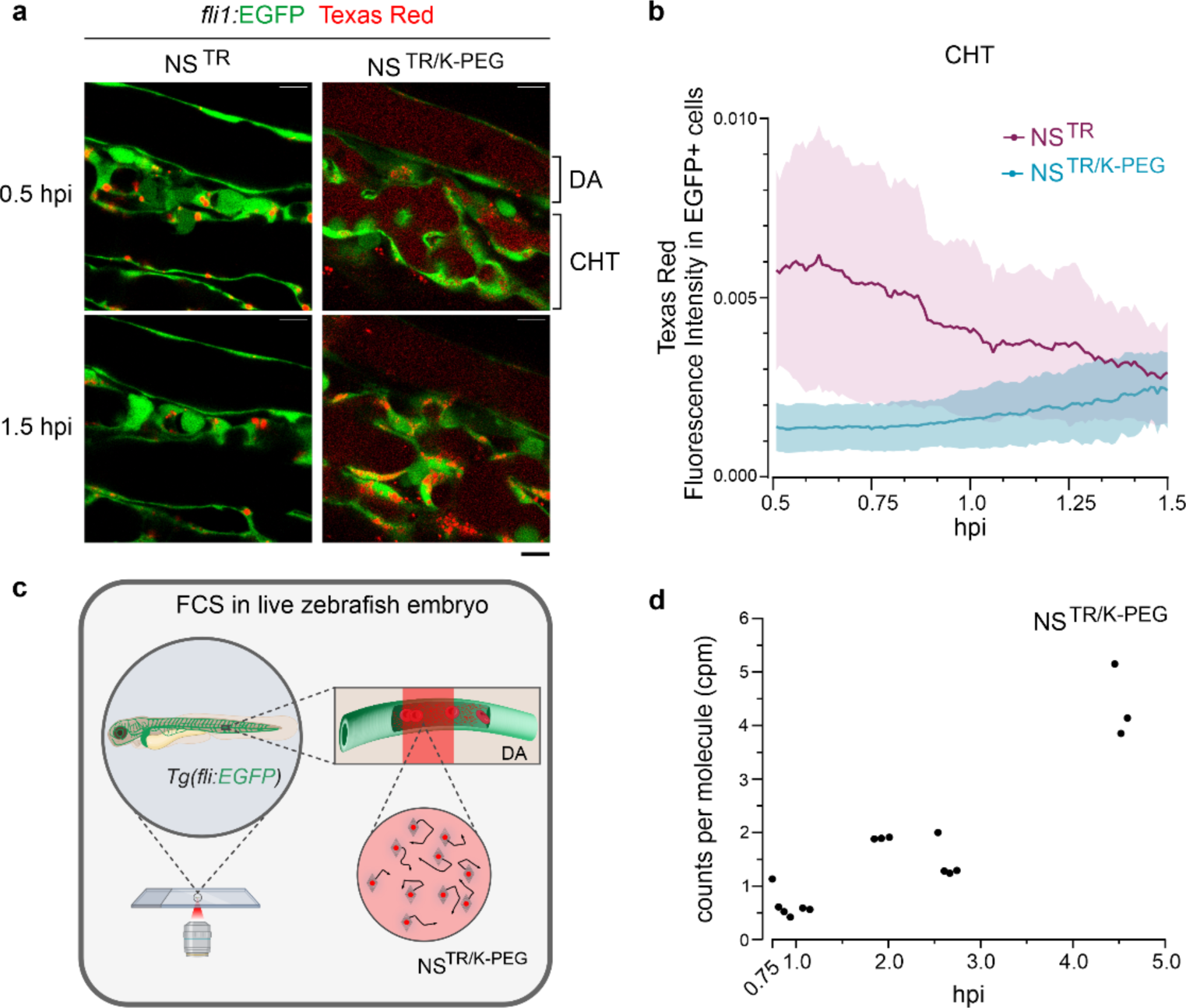
NanoSheet-cell interactions at the CHT. **a.** Lateral view images of the DA and CHT in live *Tg(fli1:EGFP)* embryos injected with NS^TR^ or NS^TR/K-PEG^. The interaction of NanoSheets with endothelial cells was recorded from 0.5 to 1.5 hpi in 30 sec intervals. n= 5 injected embryos per condition. Scale bar 10 μm. **b.** Quantification of the NS^TR^ and NS^TR/K-PEG^ levels in EGFP^+^ cells based on Texas Red mean fluorescence intensity values from 0.5 hpi to 1.5 hpi. Values presented as mean ± SEM. **c.** Schematic of the *in vivo* FCS experiment: a live *Tg(fli:EGFP)* embryo injected with NS^TR/K-PEG^ was immobilized on a glass surface with low melting point agarose, followed by FCS measurements acquisition of the Texas Red signal at the lumen of the DA. **d.** Analysis of the brightness levels of Texas Red signal originating from NS^TR/K-PEG^ in live zebrafish embryos circulation, at the indicated time points. Brightness levels Brightness (cpm: counts per molecule) was measured by FCS starting at 0.75 hpi. n= 9 technical repeats, total 20 sec traces. The increase in cpm with time observed is likely underestimated due to fluorophore quenching in the aggregates.

To analyze NanoSheet localization to the pronephric tubes of zebrafish embryos, we used the *Tg(foxj1a:EGFP)* transgenic line, as previous studies have shown the importance of kidney clearance for rectangular DNA origami in mice^8^. We observed that there was no accumulation of NanoSheets in the pronephric tubes at 4 and 24 hpi, using confocal microscopy (Extended Data Fig. 4). Together, these results suggest that the K-PEG coating does not prevent NanoSheets from accumulating in the CHT, the embryonic liver of zebrafish. However, there was a time window of one hour before we detected accumulation of the NS^TR/KPEG^ nanostructures in the CHT whereas NS^TR^ were detected shortly after injection took place. Additionally, we observed that the K-PEG coating offered long-term protection to the NanoSheets localized to the CHT.

### Dynamics of NanoSheet clearance at the CHT

After identifying CHT as the primary tissue for NS^TR^ and NS^TR/K-PEG^ accumulation, we used Airyscan imaging to dissect its role in NanoSheet clearance in *Tg(fli1:EGFP)* embryos with subcellular resolution. We focused on the time window from 0.5 hpi to 1.5 hpi, when the fluorescence intensity curves of NS^TR^ and NS^TR/KPEG^ were significantly different (Fig. 2e). We imaged the CHT and dorsal aorta (DA) of live injected embryos (Fig. 3a). Quantification of the Texas Red signal revealed that NS^TR^ accumulated and degraded faster than NS^TR/K-PEG^ in EGFP^+^ cells (Fig. 3b). In addition, NS^TR/KPEG^ were also detected in EGFP^-^ regions, indicating their interaction with non-endothelial CHT resident cells (Extended Data Fig. 5). NS^TR/K-PEG^ were detected in the circulation for the duration of the imaging, while NS^TR^ was not detected by 1.5 hpi (Extended Data Fig. 5). To further investigate the aggregation state of NS^TR/K-PEG^ in circulation, we performed FCS in the blood stream of live zebrafish embryos. We measured the fluorescence intensity fluctuations of NS^TR/K-PEG^ diffusing through a detection volume in the lumen of the DA (Fig. 3c). Brightness analyses of the objects showed that coated NanoSheets started to form aggregates in circulation at 2 hpi, which increased in size with time (Fig. 3d). Together, these data suggest that the endothelial cells within the CHT are the primary cell-type that interacts with the NanoSheets. However, we observed that the coating affected the clearance dynamics and that NS^TR/K-PEG^ were able to interact with cells within the CHT other than the endothelial cells, unlike NS^TR^. Further, NS^TR/K-PEG^, but not NS^TR^, were detected in circulation long after the microinjections.

### Whole embryo transcriptome analysis of NanoSheet- interacting cells

To identify the cell types interacting with the NanoSheets at the whole organism level with single-cell resolution, we performed single-cell RNA sequencing (scRNAseq) on embryos injected with NS^TR^ or NS^TR/K-PEG^. Embryos were injected at 2 dpf and dissociated into single cell suspensions at 4 hpi (Fig4a). Texas Red^+^ cells were sorted by flow cytometry, for both NS^TR^ and NS^TR/K-PEG^ injected embryos (Fig. 4b and Extended Data Fig. 6). We obtained 1032 Texas Red^+^ cells from NS^TR^ injected embryos with a median of 1908 genes per cell, and 1751 Texas Red^+^ cells from NS^TR/K-PEG^ injected embryos with a median of 2076 genes per cell. We used UMAP dimensionality reduction for visualization of the clusters. Based on previously annotated data ^26,27^, we identified ten different cell types for both conditions (Fig. 4c). Further, we confirmed the identity of the cell types by comparing the analyzed genes to known cell markers (Fig. 4d). Interestingly, 45.39% of the total number of NS^TR^-interacting cells was identified as scavenger/vascular endothelial cells, whereas this percentage was only 14.03% in the NS^TR/K-PEG^- interacting cells (Fig. 4e). The largest group of NS^TR/K-PEG^- interacting cells were identified as mesenchymal neural crest cells, constituting 30% of the total population (Fig. 4e). In addition, the percentages of cells identified as embryonic brain, red blood cells (RBC), the musculature system and muscle cells were higher in NS^TR/K-PEG^- than NS^TR^-labeled cells (Fig. 4e). The percentage of immune cells was low in both conditions at this time point (4hpi) but was more than two-fold higher for NS^TR^- that NS^TR/K-PEG^- interacting cells (Fig. 4e). Together these data indicate that the K-PEG coating allows for a wider distribution of NanoSheets at the whole organism level, which facilitates their interaction with tissues other than the CHT.

**Fig. 4:**
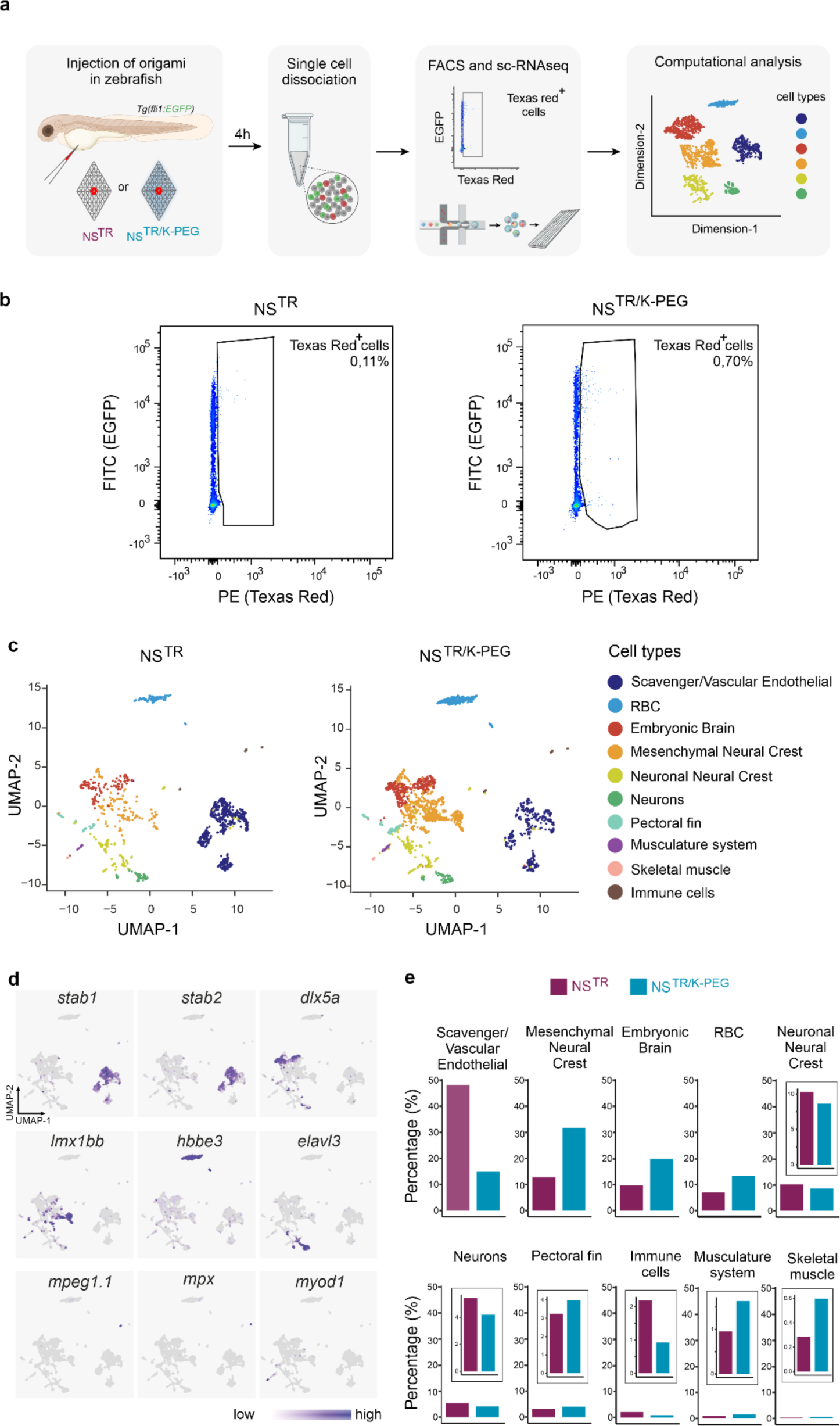
Identification of NanoSheet-labeled cell types by single-cell RNA-seq. **a.** Schematic overview of strategy to identify cells interacting with NanoSheet. *Tg(fli1:EGFP)* embryos were injected with NS^TR^ or NS^TR/K-PEG^ at 2 dpf. At 4 hpi, embryos were dissociated to single cells, Texas Red^+^ (TR^+^) cells were collected by flow cytometry and processed on a 10X platform for single-cell RNA profiling. **b.** Dot plot graphs from flow cytometry analysis representing gates for TR^+^ cells in NS^TR^ and NS^TR/K-PEG^ injected embryos. **c.** UMAP plots of 1032 TR^+^ cells from NS^TR^ and 1751 TR^+^ cells from NS^TR/K-PEG^ injected embryos. Ten different cell clusters were identified for both NS^TR^ and NS^TR/K-PEG^. **d.** UMAP plots showing the relative expression levels of selected marker genes for scavenger endothelial cells (*stab1* and *stab2*), mesenchymal neural crest (*dlx5a*), embryonic brain (*lmx1bb*), red blood cells (RBC) (*hbbe3*), neuronal neural crest (*elavl3*), immune cells (*mpeg, mpx*) and skeletal muscle (*myod1*). **e.** Percentages of identified cell-types labelled with NS^TR^ or NS^TR/K-PEG^.

### Macrophage contribution to NanoSheet clearance

It has been previously reported that liver macrophages play a significant role in the clearance dynamics of nanoparticles in zebrafish^28^. Our findings from the whole-body single-cell transcriptome analysis of zebrafish embryos at 4 hpi, indicated that few macrophages (cells expressing the macrophage specific gene marker *mpeg1*, Fig. 4e) interacted with the NanoSheets. To investigate the functional roles of macrophages within CHT in the clearance dynamics of NanoSheets at later time points, we used a transgenic zebrafish line engineered for conditionally targeted ablation of macrophages^29,30^ (Fig. 5a). Additionally, this transgenic zebrafish line expresses mCherry in macrophages which was used for their visualization, and therefore uncoated and coated NanoSheets were labeled instead with Alexa Fluor 488 and named NS^AF488^ and NS^AF488/K-PEG^, respectively (Fig. 5a). We induced macrophage ablation by treatment with metronidazole (MTZ), from 1 dpf and until the end of the experiment at 72 hpi. We observed a reduction in the numbers of macrophages detected on MTZ-treated larvae compared to controls, for the duration of the experiment (Fig. 5b,c). We injected non-macrophage-ablated or macrophage-ablated embryos with NS^AF488^ or NS^AF488/K-PEG^ nanostructures at 2 dpf and measured their fluorescence intensity by confocal microscopy at 4, 24, 48 and 72 hpi. We confirmed that in this zebrafish model, the fluorescence intensity of the NanoSheets in the CHT gradually decreased over time (Extended Data Fig. 7), consistent with the results we obtained for the *Tg(fli1:EGFP)* line (Fig. 2f). Interestingly, we observed that the fluorescence signal from NS^AF488^ was significantly higher in macrophage-ablated embryos compared to controls, at 48 hpi (Extended Data Fig. 7) and 72 hpi (Fig. 5d) but not at earlier time points (Extended Data Fig. 7). In contrast, macrophage ablation did not significantly affect the fluorescence signal from NS^AF488/K-PEG^ at these time points (Extended Data Fig. 7 and Fig. 5d). Together, these results suggest that the effects of CHT macrophages on the clearance of uncoated NanoSheets start days after the injection of the nanostructures in the blood circulation of the zebrafish. In addition, our data suggests that K-PEG coating offers protection against macrophage-mediated clearance within the CHT.

**Fig. 5:**
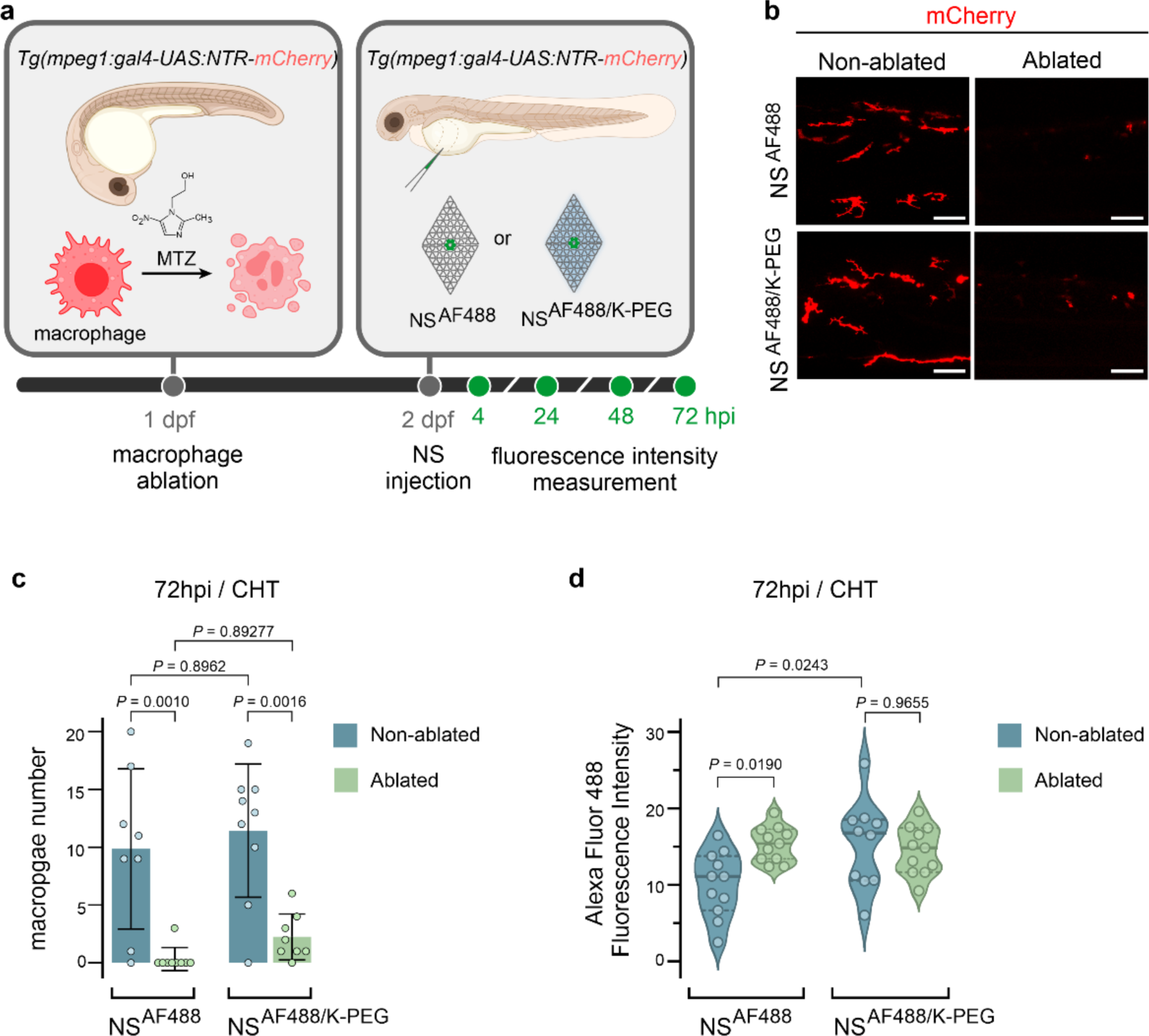
Macrophage involvement on NanoSheet clearance. **a.** Schematic strategy to evaluate the role of macrophages on NanoSheet clearance in zebrafish embryos. The *Tg(mpeg1:gal4);Tg(UAS:NTR-mCherry)* embryos were used, which expresses the nitroreductase enzyme (NTR) and the fluorescent marker mCherry under the control of the *mpeg1* promoter that drives gene expression in macrophages. mCherry labelling was used to detect macrophages. Targeted macrophage ablation is elicited by treatment with the MTZ compound, which is converted by NTR into a cytotoxic byproduct that ablates macrophages. NanoSheets used in this assay were fluorescently labelled with Alexa Fluor 488 (AF488). Embryos were treated with MTZ from 1 day post fertilization (dpf) until the end of the experiment. Embryos were intravenously injected with NS^AF488^ or NS^AF488/K-PEG^ 24 hours later (2 dpf) and imaged by confocal microscopy at the CHT region at 4, 24, 48 and 72 hpi. Different embryos were imaged at each time point. **b.** Live confocal microscopy imaging of the mCherry protein expressed in macrophages in the indicated conditions at 72 hpi. Scale bar 50 μm **c.** Bar plots of the quantifications of mCherry^+^ macrophages in CHT region at 72 hpi. Macrophage-ablated embryos were treated with MTZ from 1 dpf. Non-macrophage-ablated embryos were not treated with MTZ. Values presented as mean ± SD. n=8 embryos (NS^AF488^ non-ablated, NS^AF488/K-PEG^ ablated) or n=9 embryos (NS^AF488^-ablated, NS^AF488/K-PEG^ non-ablated) per condition. *P*-values determined by one-way ANOVA followed by Tukey’s multiple comparison test. **d.** NS^AF488^ and NS^AF488/K-PEG^ levels based on confocal AF488 mean fluorescence intensities of MTZ-treated and untreated larvae at 72 hpi. n=10 embryos (NS^AF488/K-PEG^ non-ablated) and n=11 embryos (NS^AF488^ ablated, NS^AF488^ non-ablated, NS^AF488/K-PEG^ ablated) per condition. *P*-values determined by one-way ANOVA followed by Tukey’s multiple comparison test.

## Conclusions

The DNA origami method allows for precise fabrication of nanostructures, offering control over their physical and chemical properties as well as site-specific functionalization with ligands of interest. These features position DNA origami nanostructures as promising therapeutic agents, capable of being selectively directed towards specific tissues. However, despite this potential, the evaluation of their performance within living organisms has been limited by the use of methods that lack single cell resolution. To tackle this, we propose a strategy integrating DNA nanotechnology with embryonic zebrafish models, advanced microscopy, and single-cell RNA sequencing. This approach aims to provide a comprehensive understanding of the biodistribution and clearance dynamics of DNA origami nanostructures in real-time under physiological conditions. We imaged NanoSheets within live animals and tracked their interactions with cells. Modifying NanoSheet surface chemistry with K-PEG did not affect which tissues they labeled but had a large impact on relative distribution of NanoSheets in the different tissues. Thereby, this coating approach provided a therapeutic window that prolonged NanoSheet interaction with tissues like the brain and musculature before their clearance. Furthermore, our observations highlighted the scavenger endothelial cells as the primary contributors to NanoSheet clearance, with macrophages playing a role at later stages. This work provides a roadmap for the evaluation of the performance of DNA origami nanostructures *in vivo,* essential for the development of future applications in nanomedicine.

## Acknowledgments

This work was supported by the European Research Council under the European Union’s Seventh Framework Program (617711, A.I.T.), the Swedish Research Council (2020-01856, A.I.T.), and the Knut and Alice Wallenberg Foundation (KAW 2017.0114, A.I.T.). Parts of this work were facilitated by the following core facilities: Zebrafish Core Facility at Karolinska Institutet, SciLifeLab Advanced Light Microscopy (ALM) Facility, BioImage Informatics Facility, Biomedicum Flow Cytometry Core Facility, and Biomedicum Imaging Core. Schematics in Figs. 2, 3, 4, and 5 were created with Biorender.com.

## Author contributions

C.K. designed, carried out, and analyzed the data from all experiments. L.B. performed and supervised zebrafish experiments. S.E. and H.B. provided support with light-sheet microscopy experiments. E.E. performed single-cell sequencing analyses. J.M.D. provided support with analyses and presentation of results. J.S. performed and analyzed AFM and AGE experiments and performed oxDNA simulations. C.Ka. and O.A. provided support with zebrafish experiments. S.W. performed and analyzed FCS experiments. A.I.T. directed the study. C.K. wrote the manuscript and all authors reviewed and edited the manuscript.

## Competing interests

The authors declare no competing interests.

## Methods

### NS, NS^TR^ and NS^TR/KPEG^ production

The design-specific staple strands were ordered from Integrated DNA Technologies (IDT) at a concentration of 100 μM and mixed to a final concentration of 463 nM. The scaffold strand p8064 was produced from modified m13 phage, as described before^23^. For the synthesis of the NanoSheets, 10 nM of the scaffold was mixed with 100 nM of the appropriate staple mix in 1x PBS. The folding reaction took place in a thermocycler (MJ Research PTC/225 Gradient Thermal Cycler) by following the steps: annealing by heating to 80 °C for 5 min followed by a cooling to 60 °C over 20 min, then a slower cooling to 24 °C over 14 hours, followed by removal of excess staples with 100 kDa MWCO Amicon centrifugal filters (Merck). For the NS^TR^ and NS^TR/K-PEG^ designs, ssDNA^TR^ or ssDNA^AF488^ was mixed with the nanostructures at a 1:1 ratio and annealed in a thermocycler by heating to 37 °C for 1 h, cooling to 22 °C at 0.1 °C per min, incubating at 22 °C for 14 h and cooling to 4 °C at 0.1 °C per min. Excess ssDNA^TR^ (IDT) or ssDNA^AF488^ (IDT) was removed using 100 kDa MWCO Amicon centrifugal filters (Merck). To produce the NS^TR/K-PEG^, NS^TR^ were mixed with oligolysine-PEG (K_10_-PEG_5K_, Alamanda Polymers) at a 1:1 ratio between the amines of lysines in K_10_-PEG_5K_ and the phosphates in DNA and incubated at RT for 30 min ^24^. The removal of the K_10_-PEG_5K_ coating was performed by treating the samples with chondroitin sulfate (Sigma-Aldrich) in 400x excess to the number of amines for 1 hour at 37 °C.

### AFM imaging

An epoxy adhesive was used to glue a disc of mica to the centre of a microscope slide. Using Reprorubber, a 3 cm high plastic ring was attached encircling the mica in order to form a chamber for imaging in liquid. NS were imaged by first diluting to 1 nM in TE-Mg buffer (5 mM Tris base, 1 mM EDTA, 10 mM MgCl_2_, pH 8.0) and pipetting 10 µL of the diluted nanostructures onto freshly-cleaved mica. After 30 s, 4 µL of 5 mM NiSO_4_ was added and left to incubate at room temperature for another 4.5 min. Unattached nanostructures were removed by washing the mica with 1.0 mL of 0.1 µm-filtered TE-Mg buffer, and the chamber was then filled with 1.5 mL of filtered TE-Mg buffer prior to imaging. The imaging was performed in TE-Mg buffer using a JPK Instruments NanoWizard 3 Ultra atomic force microscope set to alternating contact (AC) mode using a Bruker AC40 cantilever.

### Agarose gel electrophoresis

After folding, NanoSheets were analyzed by running samples on 2% agarose (Thermo Scientific TopVision Agarose) gels (0.5x TBE buffer, 10 mM MgCl_2_ and 0.5 mg/ mL ethidium or 1x SYBR Safe DNA stain) in an ice bath, for 4h at 75V. The gels were imaged in the ImageQuant LAS 4000 system (GE Healthcare).

### oxDNA simulation

NS were designed using the vHelix program, a plugin for Autodesk Maya. The designs were converted to the oxDNA format using the tacoxDNA web site (http://tacoxdna.sissa.it/) and analyzed using oxDNA coarse-grained modelling through the browser-based platform at https://oxdna.org/. Simulations of the nanostructures were carried out on the oxdna.org web server with the following parameters: 37°C, a salt parameter of 1, 1 x 10^8^ timesteps with a dt value of 0.0001, and a preliminary relaxation step with the default parameters. Visualizations and videos of the simulations were made using the oxView tool (https://sulcgroup.github.io/oxdna-viewer/).

### *In vitro* FCS (sample preparation + measurements acquisition)

ssDNA^TR^, NS^TR^ and NS^TR/K-PEG^ samples were prepared at a final concentration of 10 nM in 1x PBS. A droplet of 2.5 μL was transferred on a glass coverslip (#1.5 glass bottom dishes, MatTek Corporation). FCS measurements were performed on a Zeiss 980 confocal laser scanning microscope equipped for FCS, with a Zeiss water immersion objective and C-Apochromat 40×/1.2 NA. All samples were excited at 561 nm and fluorescence emission was collected at 570-694 nm. The FCS detection volume was calibrated using Alexa 568 (the diffusion coefficient of Alexa 568 was estimated to D=350 μm^2^/s by comparing with Alexa 488, D=414 μm^2^/s,^31^ using 488 nm excitation in both cases, yielding ω=0.25 μm and V=0.42 fL. The measurement on TR-ssDNA yielded τ_D_=115 μs corresponding to D_primer_ = 134 μm^2^/s. Measurement on NS^TR^ yielded τ_D_=2.30 ± 0.05 ms and measurement on NS^TR/KPEG^ yielded τ_D_ = 2.3 ± 0.25 ms, both corresponding to D=6.7 μm ^2^/s.

### Zebrafish microinjections and macrophage ablation

Zebrafish were housed in self-cleaning 3.5 l tanks with a density of 5 fish per liter in a centralized recirculatory aquatic system (Tecniplast). Basic water parameters were continuously surveilled and automatically adjusted to a temperature of 28 °C; conductivity 1200 µS/cm, pH 7.5. Other chemical water parameters were checked minimum monthly. The lightning scheme was 14 hours light/10 hours dark with a 20 min dawn and dusk period. Any animals are imported to a physically separate quarantine unit from which only surface-disinfected eggs are transferred to the breeding colony barrier. Health monitoring was done through Charles River according to the FELASA-AALAS guidelines^32^. Mycobacterium chelonae has been detected in sludge samples, ZfPV-1 in sentinel fish. Historically, Pseudoloma neurophilia had been detected in sentinels. Zebrafish embryos were staged according to Kimmel et al.^33^. All husbandry procedures are defined in SOPs which are available together with the latest health monitoring reports on request. All experiments were performed on animals younger than 5 days and no ethical permit was required according to 2010/63/EU. Previously established zebrafish transgenic lines were used in this study; *Tg(fli1:EGFP)^y1^* ^34^, *Tg(foxj1a:EGFP)*^35,36^ and *Tg(mpeg1:GAL4);Tg(UAS:NTR-mCherry)*^29,30^.

For injection into the blood stream, 2 dpf embryos were manually dechorionated and anesthetized with MS-222 (Merck). 4 nl of the sample were delivered into the common cardinal vein using microcapillaries (TW100-4, World Precision Instruments), which were pulled using a Sutter P1000 needle puller.

The samples for injections were prepared at a final concentration of 100 nM in 1x PBS for the control (NS mixed with ssDNA^TR^ in 1:1 ratio), NS^TR^ and NS^TR/K-PEG^. The total blood volume for a 2 dpf zebrafish is 60–89 nl^37^ and the estimated concentration of the injected nanostructures in our assays is aproximately 4-6 nM.

For macrophage ablation double transgenic embryos *Tg(mpeg1:GAL4);Tg(UAS:NTR-mCherry)* were exposed to 10 mM MTZ (Sigma-Aldrich) diluted in 1% DMSO (VWR) in anE3 medium supplemented with 30 mg/ml 1-phenyl-2-thiourea (PTU, Acros Organics) from 24 hpf. For the assigned ablated animal groups, medium was replaced daily with freshly prepared MTZ solution up until the last imaging time point at 72 hours post injection (hpi). The quantification of macrophages was based on a scoring system with cells being categorized in three groups (type1: macrophages with long projections, type 2: macrophages with size ³10 μm in diameter, type 3 with size 10 μm in diameter and signs of morphological cell death). Type 1 and type 2 cells were included in the quantifications.

### Light Sheet imaging

*Tg(fli1: EGFP)* embryos (48 hpf) were anesthetized in 0.005% tricaine (MS222) and mounted in 1% low melting point agarose (Thermo Scientific) in a glass capillary (50 µl, BRAND GmbH) directly after they were injected with control, NS^TR^ or NS^TR/K-PEG^. The agarose cylinder was extruded into the sample chamber of a Light Sheet Z.1 microscope (Carl Zeiss, Germany) containing egg water solution (E3) at 28.5°C. The 488 and 561 nm laser lines were used to excite fluorescence and images were acquired using a water dipping 10X detection objective (W-Plan-APOCHROMAT-0.5NA) and dual side 5X illumination objectives (LSFM-5X/0.1NA). Samples were illuminated from two sides and tiled (1 x 3), Z-stacks were acquired every 4 minutes for 4 hours. The z-stacks were max intensity projected and stitched in Zen Blue 3.6 (Carl Zeiss, Germany). When required, a drift correction was applied to time-lapse images using the “Linear Stack Alignment with SIFT multichannel” plugin in Fiji. The maximum intensity projections of the *z-*stacks over time were analysed with Fiji v2.14.0^38,39^. The average intensity of control, NS^TR^ or NS^TR/K-PEG^ was measured within a 10*10 μm rectangular area in the centre of the lumen of dorsal aorta (DA) for the slice of each timepoint. This measurement was repeated in three different sites along the DA, in each embryo. All the displayed images were acquired from the same experiments and their contrast values were adjusted for visualization purposes.

### *In vivo* confocal imaging

Injected embryos were collected at different time points, anesthetized, and mounted in 1% methylcellulose (Sigma). The confocal images were acquired with a Leica TCS SP8 microscope and the LAS X software (v. 3.5.5.19976). The CHT of the animals was scanned with a ×40 water-immersion objective and the maximum intensity projections of the *z-*stacks were analysed using Fiji v2.14.0^38,39^. All the displayed images were acquired from the same experiments and their contrast values were adjusted for visualization purposes.

### *In vivo* Airyscan imaging

*Tg(fli1:EGFP)* embryos (48hpf) were injected with NS^TR^ or NS^TR/K-PEG^. Thirty minutes post injections one successfully injected embryo per group was anesthetized in 0.005 % tricaine (MS222) and mounted on a lateral position in 1% low melting point agarose (Thermo Scientific) on a cover-glass bottomed petri-dish (MatTek). Once the agarose had polymerised, the dish was filled with egg water solution (E3).

The zebrafish caudal hematopoietic tissue (CHT) was imaged on a Zeiss LSM980 Airyscan using a C-Apochromat 40x/1.2 NA water immersion objective equilibrated at 28.5 °C. Fluorescence was excited using the 488 and 561 laser lines and single-plane 2 color images were acquired every 30 seconds for 1 hour in the Airscan SR-4Y mode. Following the time-lapse acquisition, a Z-stack covering approximately 40 µm was acquired. Three dimensional reconstructions were made using Imaris v 9.7.0. Image analysis of the image intensities was performed using CellProfiler 4.2.6^40^. For segmentation of the endothelial cell regions of the image, the green channel (EGFP is expressed under the control of the promoter for the endothelial marker Fli1a) was filtered using a median filter (filter radius 30 pixel) and then segmented using the Otsu threshold method within the Identify Primary Object module. The achieved objects were expanded by 6 pixels to ensure coverage of small intracellular vesicles. The mean intensities of the red channel (DNA origami) were then measured within the expanded EGFP^+^ regions. All the displayed images were acquired from the same experiments and their contrast values were adjusted for visualization purposes.

### *In vivo* FCS measurements acquisition

*Tg(fli1:EGFP)* zebrafish embryos (48 hpf) were injected with NS^TR/K-PEG^. A successfully injected embryo was anesthetized and mounted as described above. FCS measurements were performed on a Zeiss 980 confocal laser scanning microscope equipped for FCS, with a Zeiss water immersion objective and C-Apochromat 40×/1.2 NA and equilibrated at 28.5 °C. Fluorescence of NS^TR/KPEG^ was excited using the 561 nm laser line at the lumen of the dorsal aorta above the CHT. Each FCS measurement point was 3 minutes (9x20 sec), and measurements were recorded during 224 min.

### Statistical analysis

Embryos were randomly assigned to the control, NS^TR^, NS^TR/K-PEG^, NS^TR^ injection and MTZ treatment groups. No statistical methods were used to predetermine sample size. Data collection and analysis were not performed blind to the conditions of the experiments. Graphs show all individual data points. Every embryo was injected with freshly prepared NanoSheets. For all experiments performed only successfully injected embryos were selected.

For light sheet and Airyscan imaging and for FCS analysis, several embryos were injected, one successfully injected embryo was selected and imaged in one experimental session at a time. One-way ANOVA with Tukey’s multiple comparisons test was performed for the analysis of multiple groups. Statistical analysis and graphical representation of the data were performed with GraphPad Prism 10.0.3.

### Single-cell RNA-seq preparation

For single-cell RNA sequencing of TS^+^ cells, dissociation of embryos (48 hpf) into a single cell suspension was performed as previously described^41^ with minor modifications. Four hours post injections, around 200 *Tg(fli1:EGFP)* embryos were euthanized in 0.02% tricaine and washed twice with 1X PBS. The tissue was dissociated with harsh pipetting in 0.25% trypsin-EDTA (GIBCO) and 100mg/mL collagenase (Sigma-Aldrich) for 3 min at 30°C, followed by resuspension in DMEM-10% FBS. The cells were then pelleted at 700xg for 5 min, washed once in 1X PBS, filtered through a 40 μm cell strainer (MERCK) and stained with 1mM DAPI solution (Abcam). TS^+^ cells were FACS sorted on a FACSARIA III with BD FACSDIVA software v. 9.0.1 (BD Biosciences).

scRNAseq libraries were generated from the sorted cells using 10X Genomics Chromium Single Cell 3ʹ Reagent Kits v3.1 per manufacturer’s instructions, targeting 8000 cells. The cDNA libraries were amplified with 14 PCR cycles and sequenced on using NextSeq 500/550 High Output Kit v2.5 (150 Cycles) on NextSeq 550 platform (Illumina), with a read depth of at least 25M reads per cell.

### Single- cell RNA-seq analysis

The sequencing data was processed and analyzed using the 10X Cell Ranger pipeline (v. 7.1.0)^42^ with the Danio rerio (GRCz11.105) genome and Seurat software package (v. 4.3.0)^43^ for R (v. 4.1.3). Data was filtered to remove cells with low gene count (<200), high read count (>75’000), and high percentage mitochondrial genes (>20%). The data was normalized (NormalizeData) and scaled (ScaleData) for the 2000 most variable features (FindVariableFeatures). Subsequently, we performed dimensionality reduction of the data to 100 principal components, from which the top 50 was used to generate UMAP and tSNE visualizations. The samples were integrated to allow for better comparison (FindIntegrationAchors, IntegrateData), prior to clustering using k=20 for the k-nearest neighbor algorithm and resolution 1.2 (FindNeighbours, FindClusters).

To annotate the clusters with their predicted cell type we analyzed the marker genes for each cluster (FindAllMarkers) using Wilcox Rank Sum test and compared these to known cell markers (Fig. 4). In addition, we used an annotated cell atlas from Zebrahub^26^ (2 dpf, v. 1) (https://doi.org/10.1101/2023.03.06.531398) together with the scPred package (v. 1.9.2)^44^ for training an algorithm using supervised learning, which was then used to annotate our dataset. To annotate cranial neural crest-derived cells (CNCCs) we compared previously published and annotated markers^27^ to marker genes in clusters of progenitor cells. Taken together, the predictions from the different methods all contributed to the final cell type annotation.

## Code availability

Code used for analysis of RNA-seq data is available at https://github.com/TeixeiraLab/Kolonelou-et-al

## Data availability

Singe-cell RNA-seq data files are available via the ArrayExpress database at https://www.ebi.ac.uk/biostudies/arrayexpress under accession number E-MTAB-13554. Reference Danio rerio (GRCz11.105) used for gene mapping in Cell Ranger (v. 7.1.0) (https://www.ensembl.org/Danio_rerio/Info/Index). Annotated single-cell RNA-seq atlas of zebrafish at 2 dpf (https://zebrahub.ds.czbiohub.org/data).

---

**Extended Data Fig. 1:**
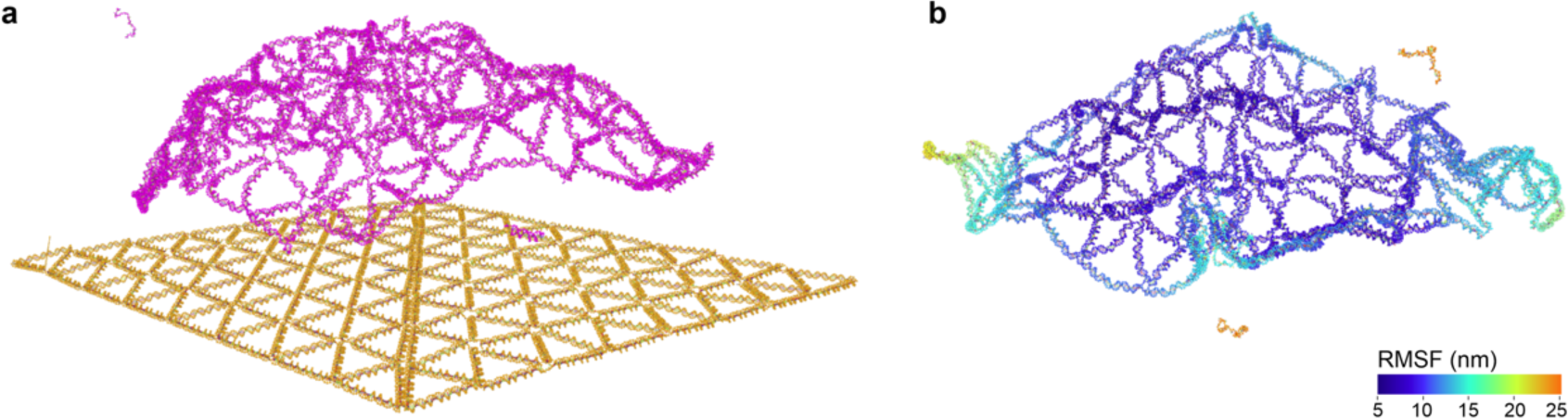
Simulation analysis of NS. **a.** Comparison between the schematic illustration of the NS design (yellow) and a simulation snapshot of NS structure generated by the oxDNA software (https://oxdna.org/) (pink). **b.** Rigidity analysis of NS with the oxDNA coarse-grained modelling software, showing a flexible structure. RMSF: root mean square fluctuation.

**Extended Data Fig. 2:**
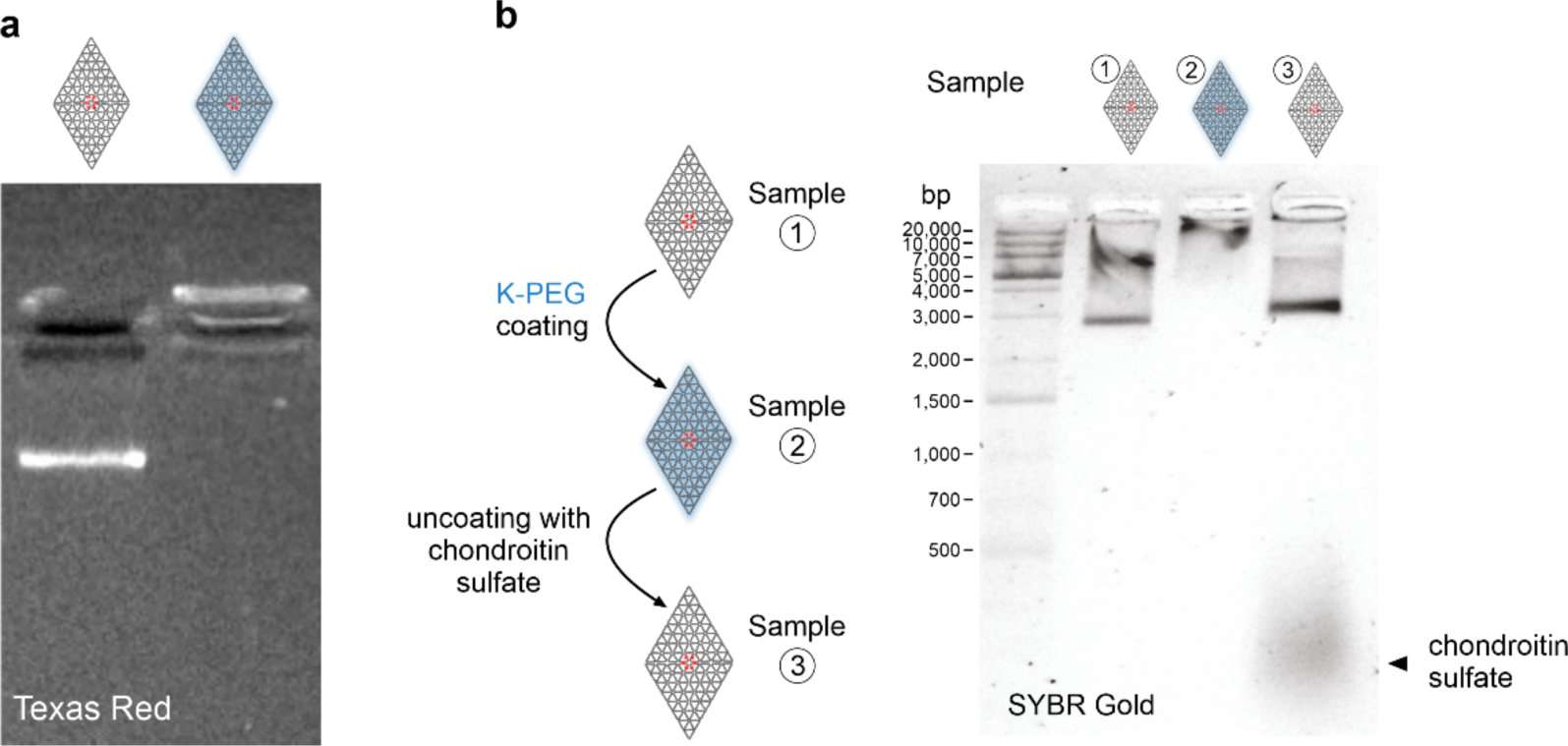
Characterization of NS^TR^ and NS^TR/K-PEG^. **a.** Analysis of NS^TR^ and NS^TR/K-PEG^ by agarose gel electrophoresis. The gel was imaged under dsCherry channel to detect the Texas Red signal. **b.** Characterization of the coating strategy of NS by agarose gel electrophoresis. Coating of NS with K-PEG sequesters the negative charges associated with the NS, resulting in a poor migration of the structures in an agarose gel electrophoresis. Treatment with the negatively charged sulfated glycosaminoglycan chondroitin sulfate electrostatically sequesters the positively charged K-PEG and fully removes K-PEG from NS^TR/K-PEG^ and re-establishes the migration pattern of NS structure in an agarose gel electrophoresis.

**Extended Data Fig. 3:**
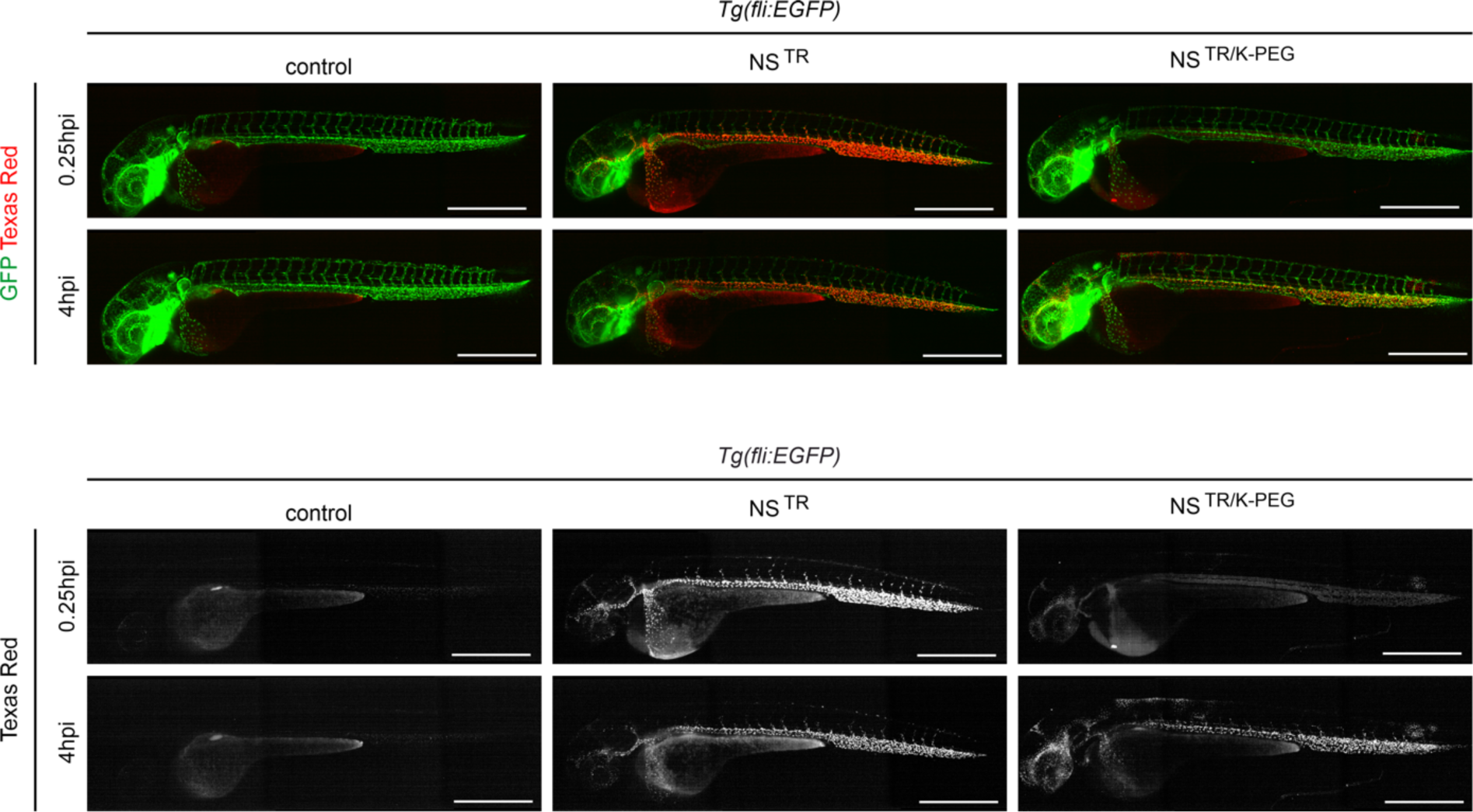
Light-sheet fluorescence microscopy images of live *Tg(fli1:EGFP)* embryos injected with control, NS^TR^ and NS^TR/K-PEG^. Biodistribution profiles of control, NS^TR^ and NS^TR/K-PEG^ at 0.25 and 4 hpi. Scale bar 500 μm.

**Extended Data Fig. 4:**
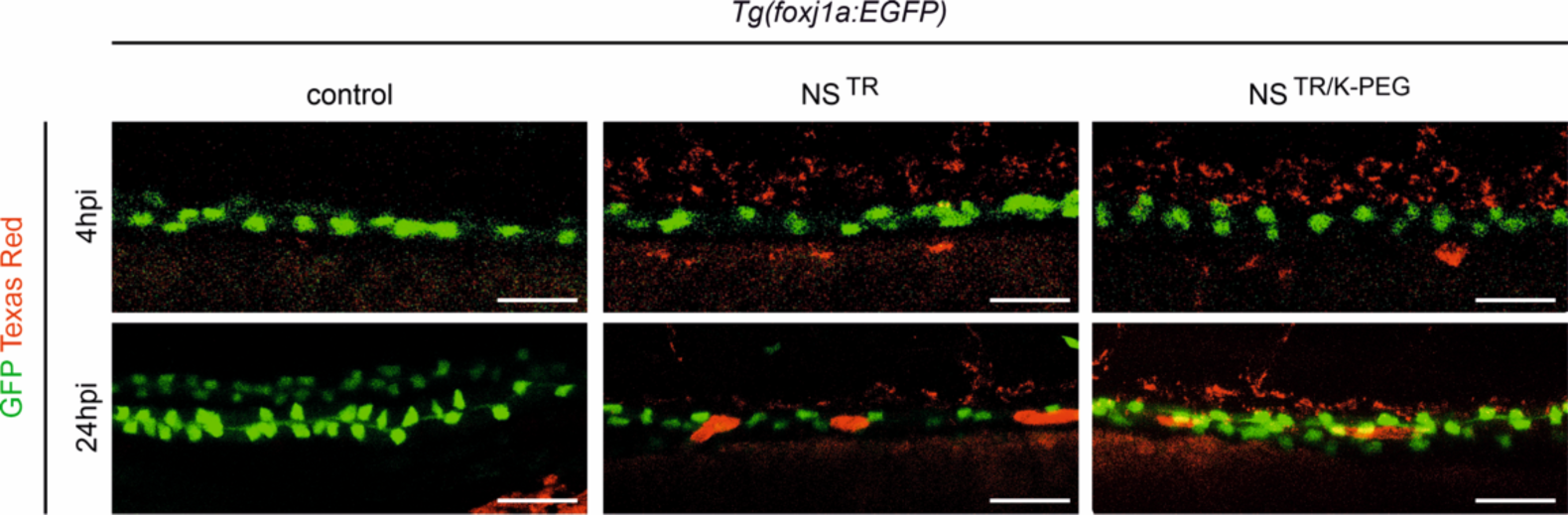
*In vivo* confocal imaging of *Tg(foxj1a:EGFP)* embryos injected with control, NS^TR^ and NS^TR/K-PEG^. Imaging of Texas red signal in the pronephric tubes (green) in control, NS^TR^ and NS^TR/K-PEG^ at 4 and 24 hpi. Scale bar 50 μm.

**Extended Data Fig. 5:**
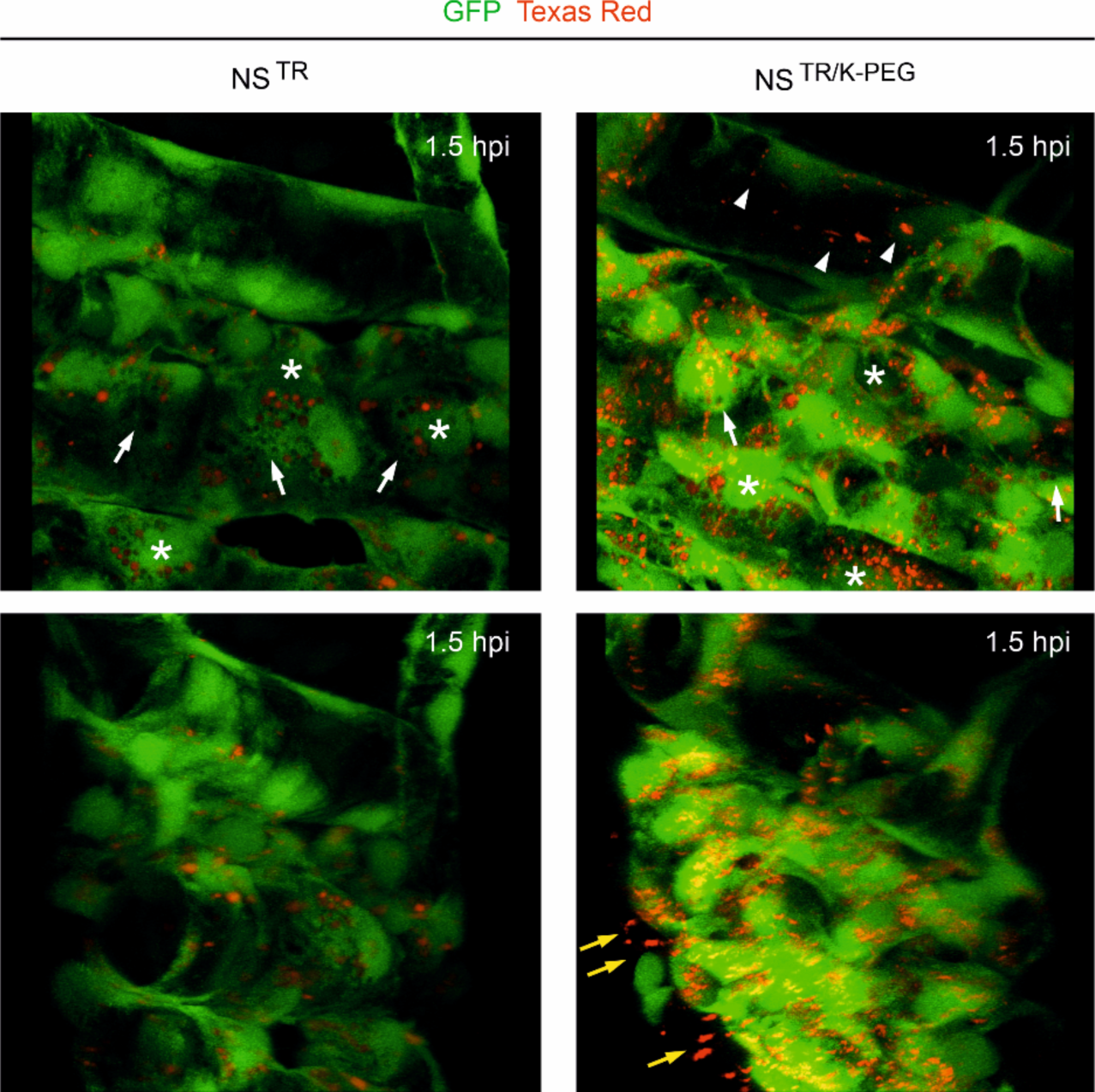
K-PEG coating prolongs circulation times of NanoSheets and promotes interactions with GFP^-^ cell-types at the CHT. Frames taken from 3D rendering of z-stack images of CHT and DA in *Tg(fli1: EGFP)* at 1.5 hpi, showing vesicles with Texas Red signal (asterisks), empty vesicles (white arrows), NS^TR/K-PEG^ in circulation (white arrowheads) and NS^TR/K-PEG^ interacting with GFP^-^ cells (yellow arrows). Embryos were injected with NS^TR^ or NS^TR/K-PEG^. Injection of each embryo was performed as an independent experiment.

**Extended Data Fig. 6:**
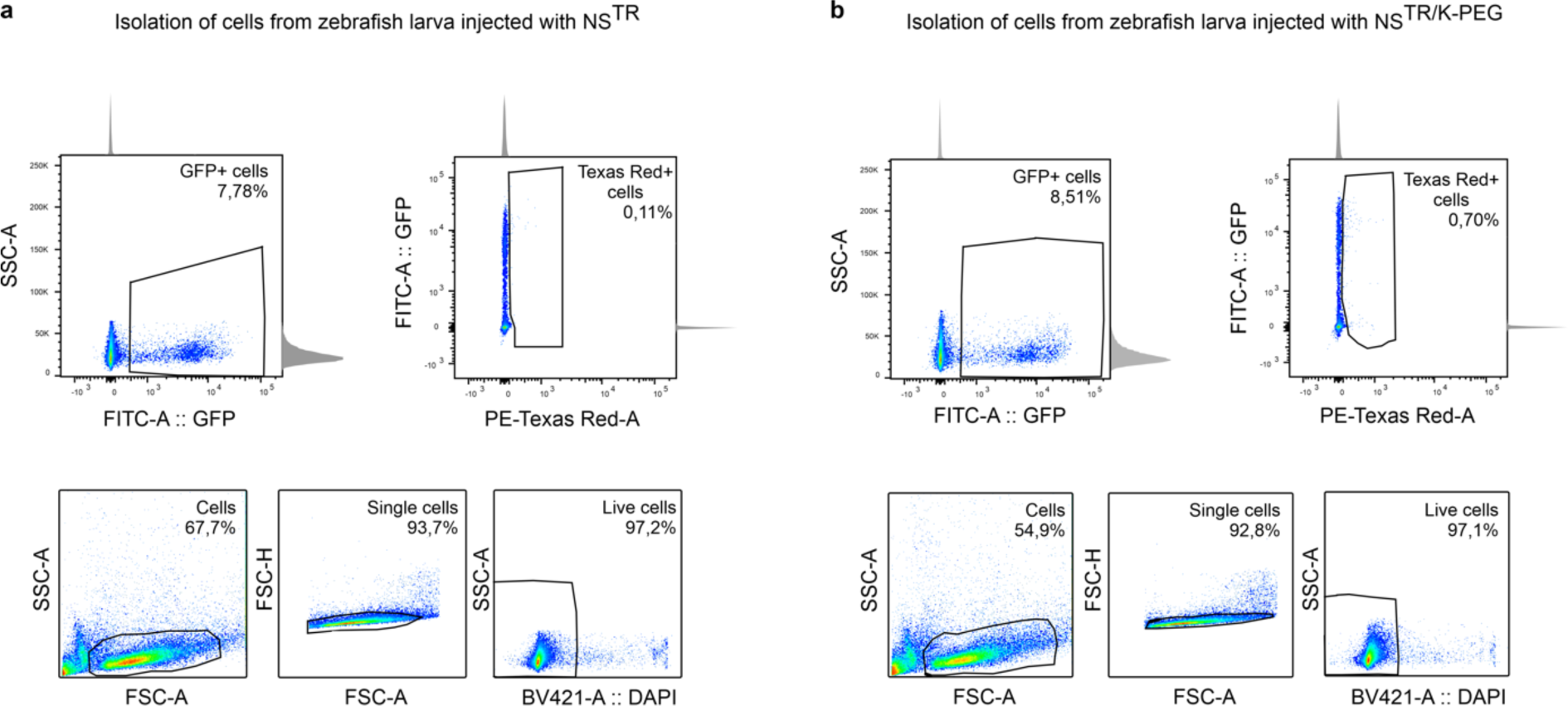
Isolation of Texas Red-labelled cells. FACS panels showing the strategy to isolate Texas Red-labelled cells from a cell suspension of whole zebrafish embryos injected at 2 dpf with NS^TR^ (a) or NS^TR/K-PEG^ (b).

**Extended Data Fig. 7:**
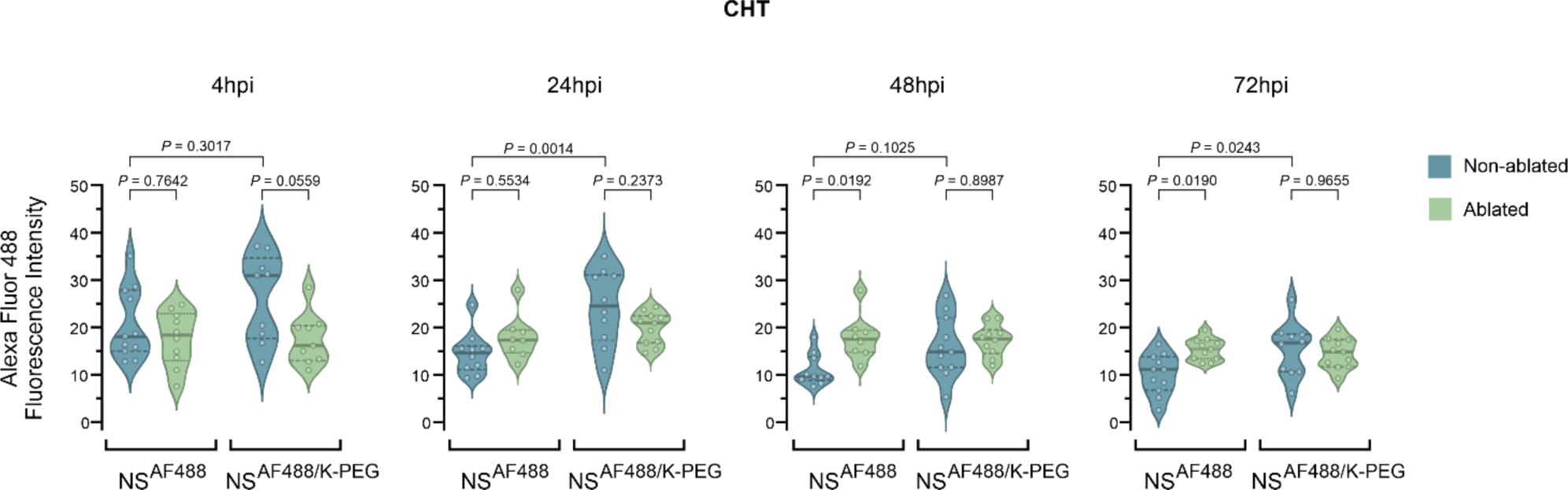
Analysis of NS^AF488^ and NS^AF488/K-PEG^ levels at the CHT in the macrophage ablated zebrafish model over time. Determination of control, NS^TR^ and NS^TR/K-PEG^ levels based on confocal AF488 mean fluorescence intensity values at the CHT of MTZ-treated (ablated) and untreated (non-ablated) embryos and larvae at 4, 24, 48 and 72 hpi. Number of individually injected embryos: n=8 (24 hpi NS^AF488^ ablated, 48 hpi NS^AF488^ ablated), n=9 ( 4hpi NS^AF488/K-PEG^ ablated, 4 hpi NS^AF488/K-PEG^ non-ablated), n=10 ( 4hpi NS^AF488^ ablated, 24 hpi NS^AF488^ non-ablated, 24 hpi NS^AF488/K-PEG^ non-ablated, 48 hpi NS^AF488^ non-ablated, 48hpi NS^AF488/K-PEG^ ablated, 72 hpi NS^AF488/K-PEG^ non-ablated) and n=11 (4hpi NS^AF488^ non-ablated, 24hpi NS^AF488/K-PEG^ ablated, 48 hpi NS^AF488/K-PEG^ non-ablated, 72 hpi NS^AF488^ ablated, 72 hpi NS^AF488^ non-ablated, 72 hpi NS^AF488/K-PEG^ ablated). *P*-values determined by one-way ANOVA followed by Tukey’s multiple comparison test.

## Notes

### Competing Interest Statement

The authors have declared no competing interest.

